# Early pathological signs in young *dysf^−/−^* mice are improved by halofuginone

**DOI:** 10.1101/762732

**Authors:** Hila Barzilai-Tutsch, Olga Genin, Mark Pines, Orna Halevy

## Abstract

Dysferlinopathies are a non-lethal group of late-onset muscular dystrophies. Here, we evaluated the fusion ability of primary myoblasts from young *dysf^−/−^* mice and the muscle histopathology prior to, and during early stages of disease onset. The ability of primary myoblasts of 5-weekold *dysf^−/−^* mice to form large myotubes was delayed compared to their wild-type counterparts, as evaluated by scanning electron microscopy. However, their fusion activity, as reflected by the presence of actin filaments connecting several cells, was enhanced by the antifibrotic drug halofuginone. Early dystrophic signs were already apparent in 4-week-old *dysf^−/−^* mice; their collagen level was double that in wild-type mice and continued to rise until 5 months of age. Continuous treatment with halofuginone from 4 weeks to 5 months of age reduced muscle fibrosis in a phosphorylated-Smad3 inhibition-related manner. Halofuginone also enhanced myofiber hypertrophy, reduced the percentage of centrally nucleated myofibers, and increased muscle performance. Together, the data suggest an inhibitory effect of halofuginone on the muscle histopathology at very early stages of dysferlinopathy, and better generation of force and muscle performance. These results offer new opportunities for early pharmaceutical treatment in dysferlinopathies with favorable outcomes at later stages of life.

## 1. Introduction

Muscular dystrophies (MDs)^1^ are genetically inherited myogenic disorders characterized by progressive muscle wasting and weakness of variable distribution and severity [1]. These symptoms are caused by cycles of myofiber degeneration–regeneration, myofiber necrosis, and the initiation of the dystrophy [2], resulting in the general pathologies of MDs, such as increased fibrosis, appearance of smaller and split myofibers, and reduced muscle mass [3–5]. The dysferlinopathies are a group of non-lethal MDs with late phenotype onset, occurring in the second or third decades of life, as opposed to other MDs such as Duchenne MD (DMD) and congenital MD (CMD) [6–8]. Although considered milder than DMD and CMD, the dysferlinopathy pathologies can severely compromise patients’ quality of life [9].

Dysferlinopathies are caused by mutations in the dysferlin gene (*DYSF*) located on chromosome 2p13. This gene encodes the ubiquitous 230-kD transmembranal dysferlin protein, located in high abundance on the sarcolemma of heart and skeletal myofibers [8,10,11]. Dysferlin has been reported to initiate the patch-repair protein complex activity that is involved in the repair of microscopic tears of the sarcolemma, and in the final stages of membrane fusion during myofiber regeneration [12,13]. Therefore, absence of dysferlin leads to delays in myofiber fusion and deficiencies in membrane resealing [3,14]. The earliest pathological signs of dysferlinopathies have been described in biopsies derived from 13-year-old patients, and current treatments focus mainly on post-disease onset [15–19]. The earliest pathological signs in mouse models for dysferlinopathies have been described in 8-week-old A/J-modified and 3-month-old SJL/J mice [14,20]. However, identification of signs of the disease at earlier ages is of the utmost importance for future development of new treatment strategies for these patients.

Halofuginone, a synthetic analogue of the alkaloid febrifugine found in the plant *Dichroa febrifuga*, has been shown to act as an antifibrotic reagent in numerous diseases [21–24]. In MDs, halofuginone has been reported to inhibit muscle fibrosis and pathology in several mouse models [25,26], including dysferlinopathies post-disease initiation, and to prevent the late outcome of the disease [27]. Moreover, halofuginone increased myofiber-attached satellite cell proliferation and myotube fusion *in vitro*, and reduced apoptosis in mouse models for DMD and dysferlinopathies, demonstrating a direct effect of halofuginone on muscle cells [28–30]. Halofuginone has recently been shown to ameliorate sarcolemmal integrity and lysosome trafficking in *dysf^−/−^* myotubes and single myofibers. Halofuginone increased synaptotagmin-7 levels and its spread across the cytosol in *dysf^−/−^* myofibers and muscle tissue, and increased its colocalization with lysosomes [31].

The mechanism of action of halofuginone is not fully understood. It has been suggested that it prevents fibroblast conversion to myofibroblasts by inhibiting Smad3 phosphorylation downstream of transforming growth factor β (TGFβ), and the entrance of phosphorylated Smad3 (P-Smad3) to the nucleus, thereby inhibiting a battery of Smad3regulated genes such as that encoding collagen type І [23,32,33]. In mouse models for MDs, inhibition of P-Smad3 by halofuginone has been reported to reduce muscle fibrosis and the pathological phenotype of dystrophic muscles [25–27]. Halofuginone has also been reported to inhibit Th17 cell differentiation via association with and reduction of prolyl-tRNA synthetase activity and activation of the amino acid starvation response [34,35].

In the present study, we show that in the *dysf^−/−^* mouse, dystrophic changes are already present at 4 weeks of age, with an initial rise in fibrosis and a delay in myotube-fusion ability. Treatment with halofuginone at this early stage delays disease onset and improves muscle histopathology and function in 5-month-old *dysf^−/−^* mice.

## 2. Materials and Methods

### 2.1. Reagents

Dulbecco’s Modified Eagle’s Medium (DMEM), sera and an antibiotic–antimycotic solution were purchased from Biological Industries (Beit-Haemek, Israel). Halofuginone bromohydrate was obtained from Akashi Therapeutics Inc. (Cambridge, MA). Hematoxylin and eosin (H&E) were purchased from Surgipath Medical Industries (Richmond, Canada).

Sirius red F3B was obtained from BDH Laboratory Supplies (Poole, UK).

### 2.2. Animals and experimental design

Male *dysf ^−/−^* [mixed 129SvJ and C57/BL/g background (Stock 006830) in which a 12kb region of *DYSF* containing the last three exons is deleted, removing the transmembrane domain] and C57/Bl/6J (Wt) mice (Jackson Laboratories, Bar Harbor, ME) were maintained as previously described [25,26]. The mice were injected intraperitoneally (i.p.) with either saline or 7.5 μg halofuginone three times a week from 4 weeks of age to 5 months of age as previously described [27]. At the end of the injection period, motor coordination was evaluated, then the mice were sacrificed and biopsies from the quadriceps, longissimus and gastrocnemius muscle were collected for further analyses [25]. All animal experiments were carried out according to the guidelines of the Volcani Center Institutional Committee for the Care and Use of Laboratory Animals (IL-234/10).

### 2.3. Motor coordination

Motor coordination and balance were evaluated with an accelerating single-station RotaRod treadmill (Med Associates Inc., St. Albans, VT) as previously described [25,26]. Briefly, the mice were placed one at a time on the rod, rotating at an initial speed of 3.0 rpm; the speed was gradually increased to 30 rpm over a period of 5 min, and the time period for which the mice stayed on the rod was recorded. RotaRod performance is highly dependent on training, therefore we performed three successive trials in each session, over two consecutive days. The performance of each mouse was measured on the second day and is presented as the mean of its best individual performances [25,26].

### 2.4. Histology and Sirius red staining

Hindlimb muscle samples were fixed with 4% (w/v) paraformaldehyde in PBS at 4°C overnight, dehydrated and embedded in paraffin as described previously [25]. Sections (5 μm) were prepared, deparaffinized and stained with Sirius red for the fibrosis evaluation. Microscopic observation and image acquisition were performed with a fluorescence microscope equipped with a DP-11 digital camera (Olympus, Hamburg, Germany). The percentage of muscle tissue occupied by fibrosis out of the total muscle tissue was calculated using FIJI (ImageJ) software (8 evenly distributed fields in 3 muscle sections were imaged per treatment).

### 2.5. Myofiber diameter analysis

The lesser myofiber diameter was determined as described previously [36,37]. Briefly, quadriceps muscle sections were stained with H&E. Ten evenly distributed fields in three to four serial sections of each muscle sample were photographed under the fluorescence microscope equipped with the DP-11 digital camera, and a total of 6000 myofibers were analyzed per treatment.

### 2.6. Cell preparation and fusion-index assay

Primary myoblasts from the hindlimb muscles of 5-week-old Wt or *dysf^−/−^* mice were prepared as described previously [29,30]. Cells were plated at 5 × 10^4^ cell/cm^2^ on 60-mm diameter Petri dishes and grown as described previously[30]. Growing myoblasts were induced to differentiate with DMEM containing 2% (v/v) horse serum for 24 h [28,30], then the medium was switched to 20% (v/v) fetal bovine serum (FBS) in DMEM for another 24 h (day 2), 96 h (day 5) and 192 h (day 8). Cells were fixed and stained for myosin heavy chain (MyHC) antibody (MF-20, diluted 1:10; Hybridoma Bank, Iowa City, IA). 488-Alexa fluor goat antirabbit IgG (diluted 1:300, Jackson Laboratories) was used as the secondary antibody. Cell nuclei were stained with 4’,6-diamidino-2-phenylindole (DAPI; 1:1000 v/v). The myoblasts were visualized under a fluorescence microscope. Myotube fusion was analyzed as previously described [27,28,38]. The number of nuclei in individual myotubes was counted for approximately 700 myotubes per day, and these were grouped into categories of cells exhibiting 2–5, 6–15, or >15 nuclei.

### 2.7. Cell preparation for scanning electron microscopy (SEM)

Primary *dysf^−/−^* myoblasts were prepared from hindlimb muscles as described in section 2.6 and plated at a density of 2 × 10^3^ cell/cm^2^ on glass coverslips (Bar Naor Ltd., Ramat Gan, Israel) in 96-well plates (Thermo Fisher Scientific, Waltham, MA), in DMEM containing 20% FBS at 37.5°C with humidified atmosphere and 5% CO_2_ in air. The cells were induced to differentiate by changing the medium to DMEM containing 2% horse serum for 24 h followed by a medium change back to 20% FBS in DMEM for 2 h. The cells were then treated, or not, with 10 nM halofuginone for another 24 h, and prepared for SEM as previously described [39]. Cells were fixed in PBS containing 2% paraformaldehyde, 2% (v/v) glutaraldehyde and 1% (w/v) sucrose for 1 h at room temperature, and treated with PBS containing 1% (v/v) osmium tetraoxide and 1% sucrose for another 1 h. The cells were then rinsed and dehydrated using increasing ethanol concentrations of 25%, 50%, 75%, 95% and 100%. The samples were dried in a critical point dryer (Bal-Tec C.P.D 030, BAL-TEC AG, Blazers, Germany) by displacing the alcohol with liquid CO_2_ and then releasing the CO_2_. The dry samples were mounted on brass blocks, sputter-coated with gold (5–7 nm) (Sputter Coater, Polaron SC 7640, Quorum Technologies Ltd., East Sussex, UK) and visualized with a JSM-5410LV scanning electron microscope (Jeol Ltd., Tokyo, Japan) under vacuum.

### 2.8. Western blot analysis

Western blot analysis was performed as described previously [40]. Briefly, equal amounts of protein were resolved by SDS-PAGE and then transferred to nitrocellulose membranes (Bio-Rad Laboratories, Hercules, CA) and reacted against the primary polyclonal antibodies goat anti-P-Smad3 and rabbit anti-Smad2/3 at 1:800 and 1:1000 dilutions, respectively (Santa Cruz Biotechnology, Dallas, TX).

### 2.9. Statistical analysis

The data were subjected to one-way analysis of variance (ANOVA) and to all-pairs Tukey–Kramer HSD test using JMP® software [41].

## 3. Results

### 3.1. Primary myoblasts from 5-week-old dysf^−/−^ mice show delayed fusion in culture

The involvement of dysferlin in membrane resealing raised the possibility of its involvement in muscle cell differentiation and myotube fusion [3,8,10,31]. Here, we addressed the differentiation and fusion abilities of *dysf^−/−^* muscle cells derived from 5-week-old mice. Primary myoblasts of Wt and *dysf^−/−^* mice were induced to differentiate, and their differentiation state was monitored by immunostaining for MyHC, a marker for muscle cell terminal differentiation. On days 2 and 5, thinner *dysf^−/−^* myotubes were observed compared to the relatively thicker Wt myotubes (Fig 1A). In addition, on day 2, despite a comparable number of nuclei in Wt and *dysf^−/−^* primary cell culture, the *dysf^−/−^* myotubes showed delayed fusion activity; these myotubes were observed mainly in areas populated by less nuclei. On day 8, although many *dysf^−/−^* myotubes were observed, they were still much thinner than those of Wt origin (Fig. 1A). The fusion index was calculated as the percentage of myotubes in each group for 2–5, 6–15 (Fig. 1B) and >15 (Fig. 1C) nuclei per myotube out of the total number of myotubes on each day. In the groups representing the smaller myotubes, the fusion index was similar in the Wt and dystrophic myotubes on all days (Fig. 1B). In contrast, the percentage of myotubes containing over 15 nuclei per myotube was zero on day 2 and significantly lower on day 8 in the *dysf^−/−^* derived cultures compared to their Wt-counterparts (Fig. 1C).

**Fig. 1.**
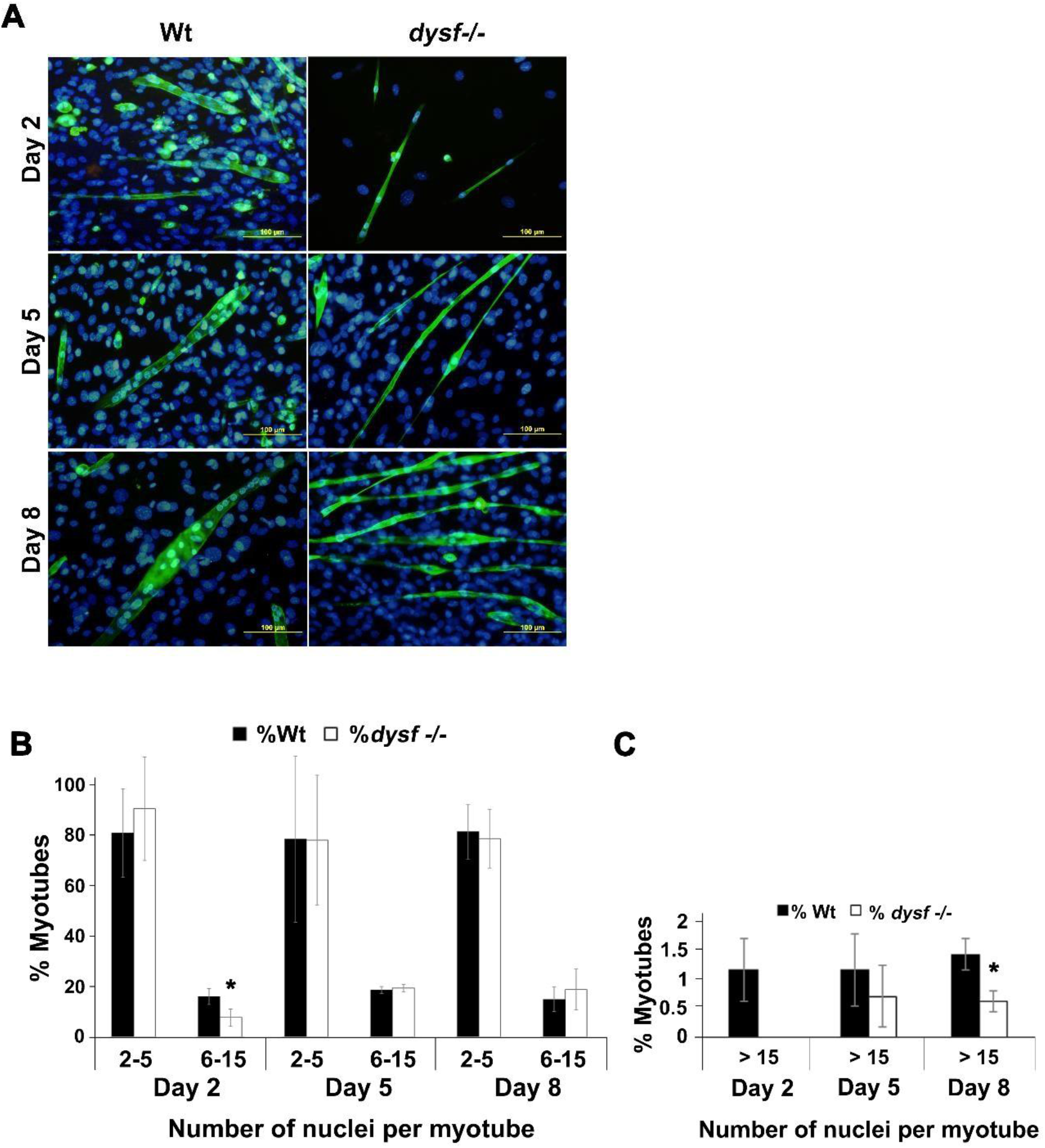
Muscle cells derived from 5-week-old *dysf^−/−^* mice show delayed differentiation and fusion abilities. (A) Myoblasts originated from Wt and *dysf^−/−^* mice were induced to differentiate with DMEM containing 2% horse serum (v/v) for 24 h, then the medium was switched to 20% FBS in DMEM for an additional 2, 5 and 8 days. The cells were fixed and stained for MyHC (green) and DAPI (blue). Note the thinner *dysf^−/−^* vs. Wt myotubes on all days. (B, C) Quantitative analysis of myotube fusion. Nuclei were counted per myotube and percentage of myotubes with 2–5 and 6–15 (B), and 15 (C) nuclei per myotube was calculated out of the total number of myotubes on each day of the experiment (~700 myotubes/day per mouse strain). Results are mean ± SE of three independent experiments. Data with asterisks differ significantly within cell origins on each day; *P* < 0.05).

### 3.2. Halofuginone has a promotive effect on dysf^−/−^ myotube fusion in vitro

The above results suggested a delay in myoblast differentiation and myotube fusion in cultures derived from *dysf^−/−^* mice at as early as 5 weeks of age. The delay in fusion might be due to impaired cell–cell connections by actin filaments—finger-like actin-based protrusions that have a role in cell movement and cell–cell fusion in many cell types, including satellite cells [42–45]. In this study, the previously reported promotive effect of halofuginone on the *dysf^−/−^* fusion index [27] was evaluated with regard to alterations in actin filaments and myotube fusion. Primary myoblasts of *dysf^−/−^* mice were induced to differentiate for 24 h as described in Fig. 1, followed by incubation with or without 10 nM halofuginone for another 24 h. The myotubes were then fixed and visualized by SEM. Under low magnification, in the non-treated myotubes, a single thin elongated myotube (white arrow in Fig. 2A) was viewed with no noticeable fusion connections to the adjacent muscle cells. However, in the halofuginone-treated myotubes, a myotube of similar size appeared to have many filaments, presumably actin filaments [44,46–48], connecting it to other myotubes (Fig. 2B, yellow arrows). Under higher magnification, many filament connections were noticed in the halofuginone-treated myotubes (Fig. 2D) compared to the non-treated myotube, which barely presented any connecting filaments (Fig. 2C).

**Fig. 2.**
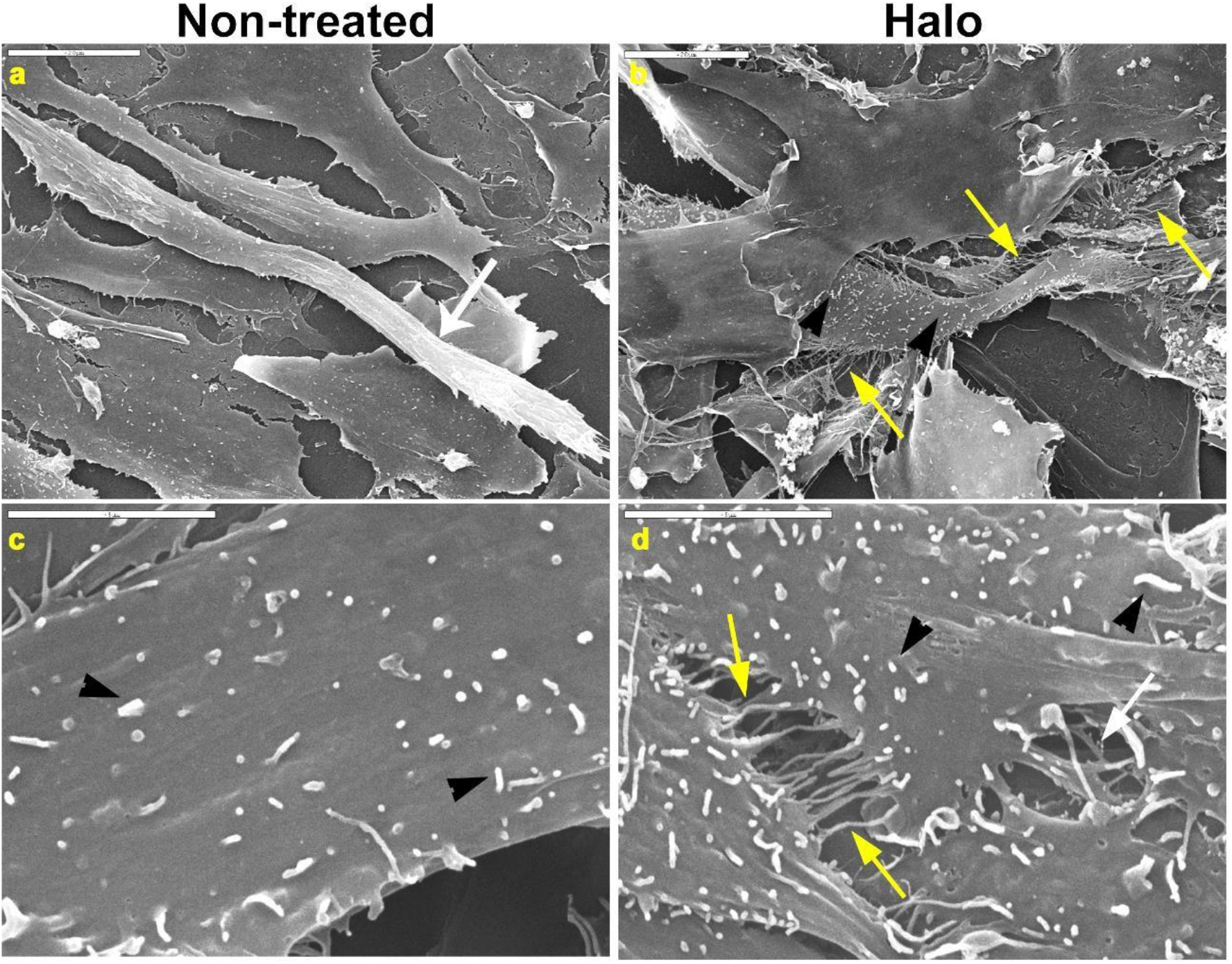
Promotive effect of halofuginone on *dysf^−/−^* myotube fusion *in vitro*. Myoblasts derived from *dysf^−/−^* mice were induced to differentiate as described in Fig. 1 and then treated, or not, with 10 nM halofuginone for 24 h. The cells were sputter-coated with gold and viewed by SEM. A myotube connected to other myotubes by filamentous protrusions (presumably actin filaments, yellow arrows) is observed in the halofuginone-treated (Halo) cells (B), whereas in the nontreated cells (A), a single thin myotube is depicted (white arrow) without any connecting filaments (magnification of X2500). At higher magnification (X10,000), fusion of several myotubes in Halo cultures (D) is observed by actin filament protrusions (yellow arrows) as opposed to hardly any filaments connecting single non-treated myotubes (C). Filipodia are noticeable on both halofuginone-treated and non-treated myotubes (black arrowheads). Bars = 20 ◻m (A, B), 5 ◻m (C, D).

### 3.3. Treatment with halofuginone decreases the levels of fibrosis and P-Smad3 in young dysf^−/−^ mice

In light of the results indicating impaired myotube fusion in cultures derived from young *dysf^−/−^* mice, and the promotive effect of halofuginone on this process, we next tested halofuginone’s effects on fibrosis in young *dysf^−/−^* mice at early stages of disease progression. Mice were treated, or not, with halofuginone from 4 weeks of age to 5 months of age (i.p. injections of either 7.5 μg halofuginone or saline, respectively, three times per week [25,26]). Hindlimb muscle sections of 4-week-old and 5-month-old Wt and non-treated or halofuginonetreated *dysf^−/−^* mice were stained with Sirius red to evaluate collagen (type I and III) content in the fibrotic tissue. Unlike in the Wt muscle sections (Fig. 3Aa), collagen staining was already notable in the myofiber fascicles of the *dysf^−/−^* muscle sections at 4 weeks of age (Fig. 3Ab). Quantitative analysis revealed a collagen content of about 0.5% in 4-week-old Wt muscle sections, whereas in the *dysf^−/−^* muscle sections, the percentage was approximately double that (Fig. 3B). At 5 months of age, in contrast to the Wt mice (Fig. 3Ac), the amount of collagen staining in the muscle sections was further increased in the non-treated *dysf^−/−^* mice (Fig. 3Ad) and was over threefold higher in the latter compared to the Wt (Fig. 3B). Treatment with halofuginone from the age of 4 weeks to 5 months reduced the collagen content in the muscle sections back to Wt levels (Fig. 3Ae). Collagen content was almost negligible in these mice and comparable to the levels in Wt mice (Fig. 3B). The reduction in fibrosis correlated with the reduction in P-Smad3 levels: western blot analysis of protein lysates derived from hindlimb muscles of 5-month-old *dysf^−/−^* mice revealed fivefold lower P-Smad3 levels in the halofuginonetreated mice than in their non-treated counterparts (Fig. 3C).

**Fig. 3.**
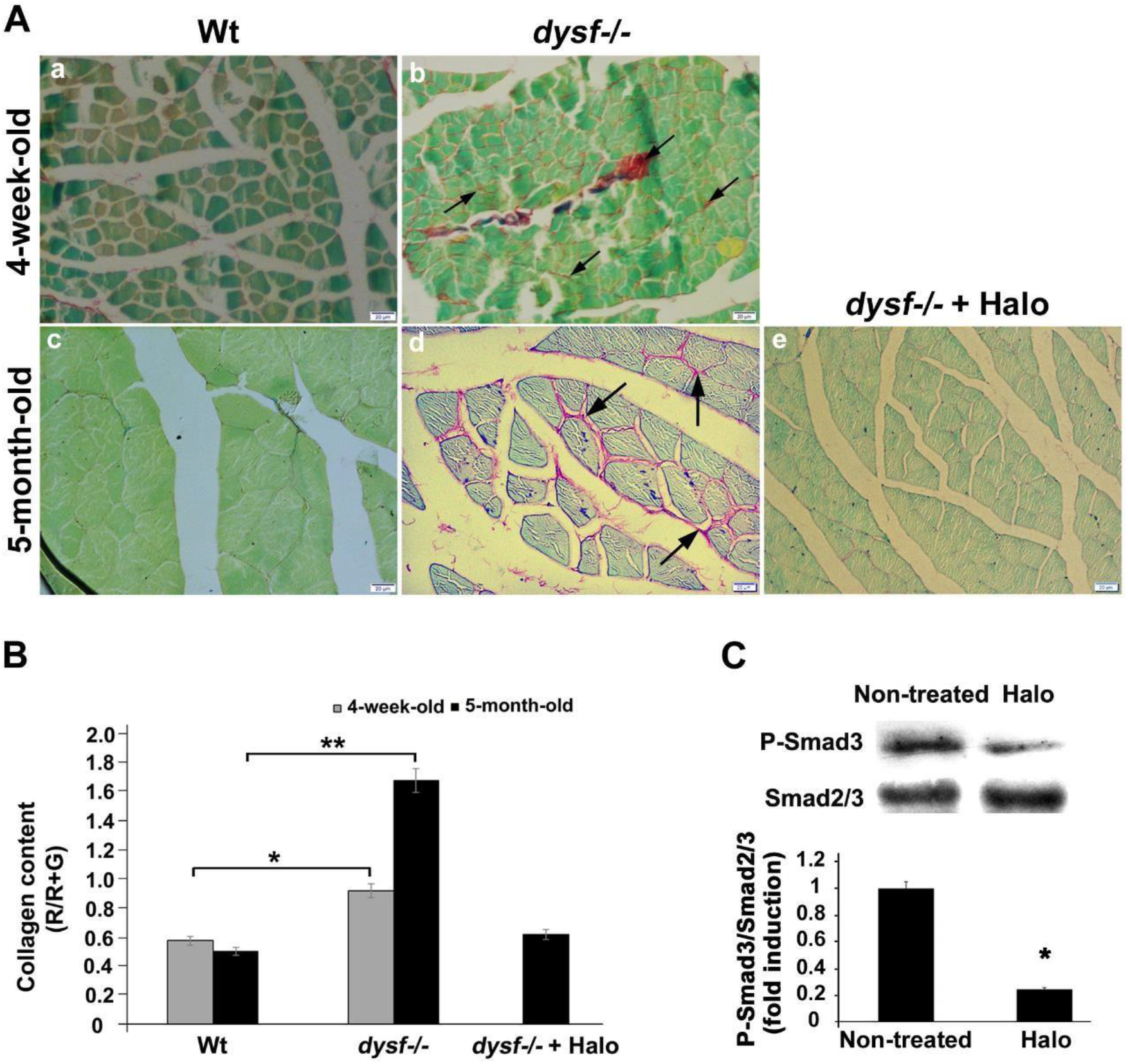
Halofuginone improves muscle histopathology of 4-week-old and 5-month-old *dysf^−/−^* mice. Saline (*dysf^−/−^*) or halofuginone (7.5 μg/mouse) was injected i.p. three times a week, from 4 weeks to 5 months of age. Non-treated Wt mice served as controls. (A) Sirius red staining of hindlimb muscle sections derived from 4-week-old (a) and 5-month-old (c) non-treated Wt mice and halofuginone-treated (*dysf^−/−^* + Halo, e) or non-treated *dysf^−/−^* (*dysf^−/−^*) mice (b, d). Note the increased collagen content between the myofibers in the *dysf^−/−^* muscle (arrows). (B) Quantitative analysis of collagen content in hindlimb muscles. Results are presented as mean ± SE (n = 3). Asterisks represent significant differences within treatments at the same age at \**P* < 0.05 and \*\**P* < 0.001. (C) Western blot analysis of the phosphorylation levels of Smad3 (P-Smad3) in protein lysates from hindlimb muscles of 5-month-old *dysf^−/−^* mice injected with halofuginone (Halo) or saline (Non-treated). Densitometry analysis was normalized to total Smad2/3 and is presented as fold-induction. Results are presented as mean ± SE (n = 3). Asterisk represents significant difference within treatments (*P* < 0.05).

### 3.4. Halofuginone improves muscle histopathology and motor coordination of 5-month-old dysf^−/−^ mice

Quadriceps muscle sections of 5-month-old *dysf^−/−^* mice, treated, or not, with halofuginone from 4 weeks to 5 months of age, were stained with H&E. Large degenerated areas with infiltrating immune cells (e.g., macrophages, yellow arrowheads in Fig. 4A) and numerous centrally nucleated myofibers (i.e., regenerated myofibers, black arrows in Fig. 4A) were observed in the muscle sections of the non-treated mice (Fig. 4A, Non-treated). In contrast, improved histopathology was demonstrated in the halofuginone-treated mice, with less inflammation and fewer myofibers with central nuclei (Fig. 4A, Halo). Quantitative analysis of 1000–1200 myofibers per mouse revealed that halofuginone significantly decreases the percentage of centrally nucleated myofibers out of the total number of myofibers (54.5 ± 0.17% and 47 ± 0.18% in non-treated and halofuginone-treated mice, respectively; n = 5; *P* < 0.0001). Myofiber diameter distribution was also evaluated in these sections. The myofiber diameters were clustered in bin intervals of 0.5 ◻m, and the percentage of myofibers in each group was calculated as the proportion of the total number of myofibers per treatment. Treatment with halofuginone resulted in a noticeable shift of the myofiber distribution curve toward the higher myofiber diameters compared to the non-treated distribution (Fig. 4B). Mean myofiber diameter was significantly higher in the halofuginone-treated mice than in their non-treated counterparts (Fig. 4C). Physiological muscle performance test on the RotaRod with halofuginone-treated or non-treated mice at the age of 5 months revealed that halofuginone markedly increases motor function and coordination: halofuginone-treated mice stayed on the RotaRod about 2.5 times longer than the non-treated mice (Fig. 4D).

**Fig. 4.**
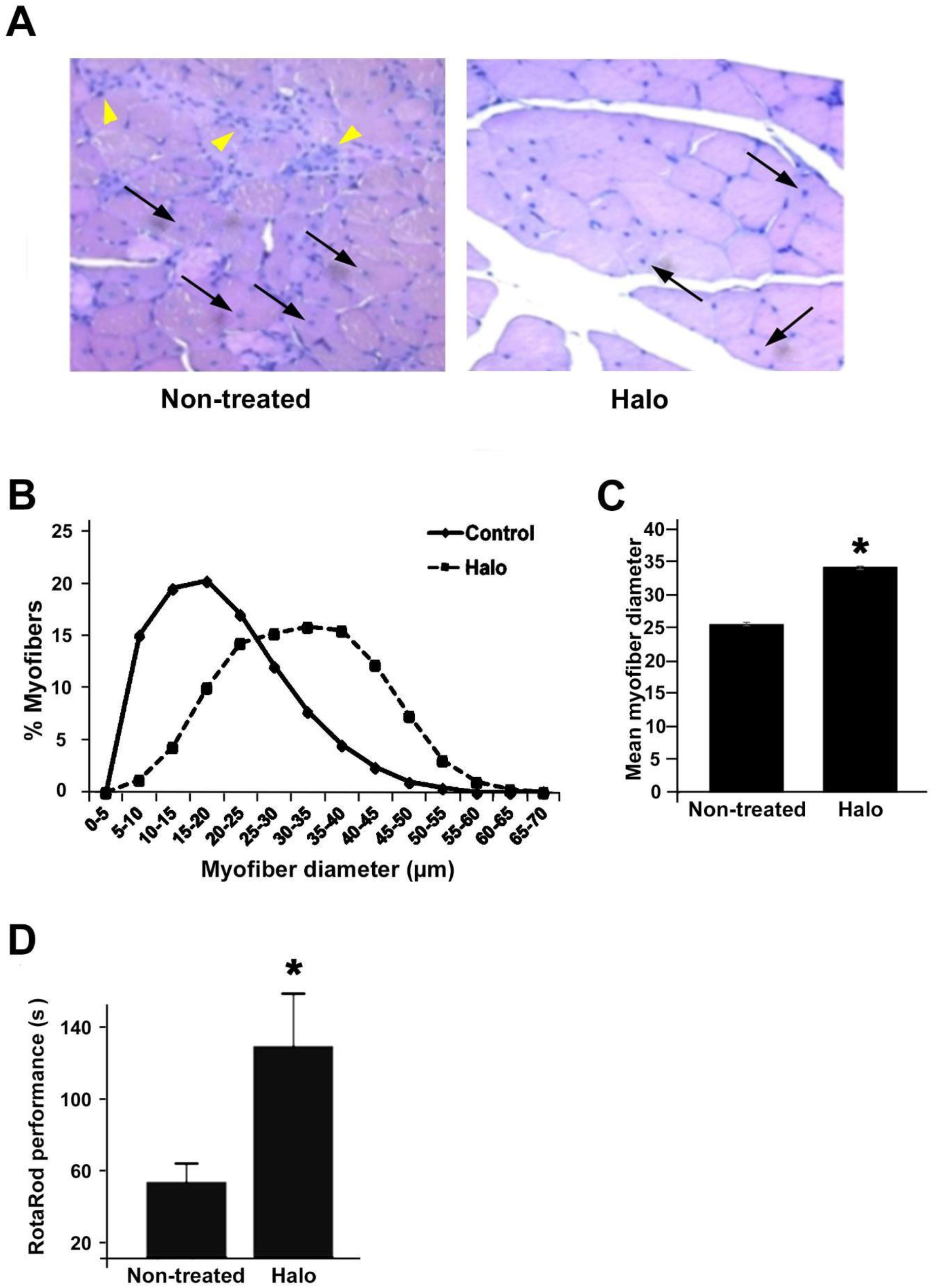
Halofuginone improves muscle histopathology and motor coordination of 5-month-old *dysf^−/−^* mice. Mice were treated, or not, with halofuginone as described in Fig. 3 and *dysf^−/−^* quadriceps muscle sections were stained with H&E. (A) Halofuginone treatment decreases the amount of myofibers with central nuclei (black arrows) and areas of macrophages infiltrating into the muscle tissue (yellow arrowheads). (B) Myofiber diameter distribution in hindlimb muscle. Myofibers were clustered in bin intervals of 0.5 μm, and results are presented as percentage of total myofibers (6000 per treatment group). (C) Mean myofiber diameter (mean ± SE). Asterisk represents significant difference between treatments (*P* < 0.05). (D) Quantitation analysis of mouse performance on the RotaRod. Results are expressed as mean ± SE of the best individual performances of the mice per treatment group (n = 5) over three trials on the second day of exercise. Asterisk represents significant difference between treatments (*P* < 0.05).

## 4. Discussion

Dysferlinopathies are considered a late-onset MD with disease manifestation during the patient’s second or even third decade of life [16,49,50]. Dysferlinopathies exhibit a milder level of fibrosis than DMD or CMD [51,52]. However, being a genetically inherited disease [53,54], early dystrophic symptoms are probably present from birth. Some dystrophic changes have been reported to appear in the distal muscles of 2-month-old *dysf^−/−^* [55] and A/J mice [14]. Nevertheless, our results, in agreement with others [56,57], demonstrate that in *dysf^−/−^* mice, fibrosis is already present at 4 weeks of age, suggesting the initiation of inflammation and cycles of myofiber degeneration–regeneration leading to fibrosis long before the appearance of the disease phenotype. The fact that fibrosis levels almost doubled from 4 weeks to 5 months of age in *dysf^−/−^* mice, together with earlier reports on pathological signs in young patients and mice [14,16,20], indicates that the disease is already actively progressing in the early stages of life.

One of the reasons for the early appearance of fibrosis is due to delayed ability of *dysf^−/−^* myoblasts to fuse and form large myotubes compared to their Wt counterparts as was observed in primary cultures derived from 5-week-old mice (Fig. 1). The delayed fusion of the myoblasts was manifested by the appearance of thinner myotubes compared to the Wt ones. One of the reasons for this phenomenon might be impairment in the myotubes’ ability to form actin filaments (Fig. 2), which are crucial for cell movement and the cell–cell fusion process [44,45,58–60]. It has also been demonstrated that under local membrane-injury protocols, dysferlin recruitment to the injury site is impaired if the formation of actin filament protrusions is inhibited [61]. Because dysferlin plays a main role in myofiber formation and regeneration [12], these results reinforce dysferlin’s role in muscle-fusion events in *dysf^−/−^* mice as young as 5 weeks of age. Halofuginone has been reported to improve the myotube fusion index of 5-week-old *dysf^−/−^* mice [27]. Here, we demonstrated the actual fusing activity under SEM: the halofuginone-treated myofibers actively fused, as observed by the many threads of actin filaments connecting several cells, whereas the non-treated myofibers presented barely any fusion activity (Fig. 2). Our evidence implies that these filaments are lamellipodia resulting from actin polymerization, causing finger-like actinbased protrusions at the muscle cells’ fusion sites [44,60,62]. This reorganization of actin cytoskeletal structures, as a result of actin or actin-regulatory protein phosphorylation, is mediated by the PI3K/Akt-signal-transduction pathway [63,64]. Halofuginone has been reported to enhance Akt phosphorylation levels in association with its promotive effect on satellite cell survival, differentiation and myotube fusion in different MD mouse models [27–30]. Therefore, it is plausible that halofuginone also increases lamellipodium formation via the PI3K/Aktsignaling pathway.

Cellular vesicles have been shown to assist in the trafficking of cell-adhesion molecules and/or fusogenic proteins to the plasma membrane [60,65]. Recently, halofuginone has been shown to rescue lysosomal trafficking to the sarcolemma in myoblasts and myofibers, probably in a synaptotagmin-7-dependent manner [31]. It may well be that the promotive effects of halofuginone on *dysf^−/−^* muscle cell fusion are also due, at least in part, to its role in the enhancement of lysosome trafficking and exocytosis during myotube-fusion processes.

We previously demonstrated halofuginone’s ability to inhibit dystrophic symptoms in *dysf^−/−^* mice at 12 months of age, when provided for short periods prior to and during the disease outburst and phenotype appearance [27]. Here, halofuginone’s ameliorative effects on muscle function and histopathology are demonstrated at the early stages of the dystrophic changes’ appearance, during the first months of life. At a very early age, halofuginone affected multiple steps in the disease’s progression. First, early treatment of *dysf^−/−^* mice with halofuginone reduced muscle histopathology in general and muscle fibrosis in particular (Fig. 3), probably by inhibiting myofibroblast differentiation to fibroblasts [23], and reducing the infiltration of inflammatory cells observed in other MDs [26,29]. Second, halofuginone increased myofiber diameter (Fig. 4) and decreased the number of centrally nucleated myofibers, in agreement with our previous reports [25,26], suggesting an ameliorative effect on the repeated degeneration–regeneration cycles and loss of myofibers. The outcome of all of these beneficial effects is better generation of force and muscle performance [66,67]. Indeed, we found a marked increase in physiological muscle motor coordination and function (Fig. 4D).

In accordance with the fibrosis reduction in the *dysf^−/−^* hindlimb muscle, Smad3 phosphorylation levels were greatly reduced by halofuginone treatment (Fig. 3). It is accepted that halofuginone’s mode of action involves the inhibition of Smad3 phosphorylation downstream of TGFβ signaling [28]. Halofuginone has also been shown to inhibit prolyl-tRNA synthetase activity in Th17 cells under conditions of amino acid starvation [34,35,68], by forming a ternary complex via halofuginone’s hydroxyl group with ATP and the enzyme, thus acting as an ATP-competitive inhibitor [69,70]. A very recent study of ours reported the requirement of halofuginone’s hydroxyl group for its promotive effect on fibrosis in mdx mice [71], raising the possibility that prolyl-tRNA synthetase inhibition by halofuginone plays a part in the latter’s effects on fibrosis in MDs, probably at early stages of the disease.

## 5. Conclusions

We demonstrate the very early appearance of dystrophic changes in *dysf^−/−^* mouse muscles; these are manifested by lower fusion index, increased degeneration–regeneration cycles and fibrosis, and attenuated levels of myofiber hypertrophy resulting in lower muscle motoric abilities. The reduced hypertrophy is probably due to delayed myoblast fusion and defects in the myotube’s ability to fuse properly, due to the absence of dysferlin. Halofuginone affects this cascade of events, and acts as an efficient reagent for the improvement of muscle cells’ abilities to fuse and form mature functioning myofibers, as well as for the amelioration of muscle histopathology and function at very early stages of disease initiation. Current treatments for dysferlinopathy focus on patients post-disease onset [18,19]. The results presented here offer new opportunities for early pharmaceutical treatment in dysferlinopathies with favorable outcomes at later stages of life; moreover, halofuginone is a likely candidate to fulfill the requirements for such a treatment.

## Acknowledgments

The authors thank Dr. Einat Zelinger for her technical assistance with the scanning electron microscopy.

## Footnotes

1 Abbreviations MD: muscular dystrophies DAPI: 4’,6-diamidino-2-phenylindole DMEM: Dulbecco’s Modified Eagle’s Medium FBS: fetal bovine serum H&E: hematoxylin and eosin MyHC: myosin heavy chain SEM: scanning electron microscopy TGF◻: transforming growth factor

